# Pharmacological inhibition of G protein-coupled receptor kinase 5 decreases high-fat diet-induced hepatic steatosis in mice

**DOI:** 10.1101/2025.06.25.661655

**Authors:** Mary E. Seramur, Sandy Sink, Tony E. Reeves, Leah C. Solberg Woods, Chia-Chi Chuang Key

## Abstract

G protein-coupled receptor kinase 5 (GRK5) is implicated in the pathogenesis of obesity in both humans and rodent models. Our previous work demonstrated that genetic deletion or pharmacological inhibition of GRK5 suppresses 3T3-L1 adipocyte differentiation. Here, we assessed the small-molecule GRK5 inhibitor, GRK5-IN-2, for its effects on metabolic tissues and therapeutic potential in a diet-induced obesity mouse model. Mice were fed a high-fat diet for 8 weeks to induce obesity, followed by continued a high-fat diet with oral administration of GRK5-IN-2 (25 or 50 mg/kg) or water vehicle, five days per week for an additional 16 weeks. GRK5-IN-2 treatment had no effect on body weight, fat/lean mass, insulin tolerance, food intake, or energy expenditure but significantly reduced hepatic triglyceride accumulation and de novo lipogenesis. A follow-up study using 25 mg/kg of GRK5-IN-2 confirmed no effect on adiposity but reduced hepatic triglycerides. GRK5-IN-2 treatment decreased expression of the lipogenic gene Acc2 while upregulating lipid utilization proteins COXIV and ACSL1 in the liver, likely contributing to lower triglyceride levels. Together, these findings suggest that GRK5 inhibition selectively modulates hepatic lipid metabolism without altering systemic metabolic parameters, highlighting GRK5 as a potential therapeutic target for fatty liver disease.

## INTRODUCTION

Obesity is a growing health concern where prevalence of overweight and obesity in the United States has increased by 50% over the past three decades and is expected to reach 80% by 2050 (1). Obesity is caused by an increase in energy intake compared to energy expenditure, where positive energy balance results in adipocytes becoming lipid-overloaded (hypertrophy) and promotes adipose tissue remodeling by increasing adipocyte number (hyperplasia) (2). The expansion of visceral adipose tissue is associated with impaired adipocyte functionality (3). This, coupled with the accumulation of ectopic lipid in peripheral organs, such as liver, contributes to the development of metabolic syndrome (4, 5).

Chronic comorbidities associated with obesity include type 2 diabetes, cardiovascular disease, and metabolic associated steatotic liver disease (MASLD), previously known as non-alcoholic fatty-liver disease or NAFLD (6, 7, 8, 9). MASLD is diagnosed when hepatic steatosis is present along with at least one of five cardiometabolic risk factors: elevated body mass index, fasting glucose, fasting triglycerides, or blood pressure, or reduced HDL-cholesterol (9, 10). Hepatic steatosis is characterized by the accumulation of lipids that exceed 5% of liver weight with no significant liver damage including inflammation or fibrosis (9, 11). Hepatic steatosis is known to present in almost every chronic liver condition (9). Although no pharmacological treatments are approved for hepatic steatosis, the condition can be reversed through sustained lifestyle interventions, including dietary changes, weight loss, and increased physical activity, which are difficult to maintain long-term (12). A meta-analysis found that weight loss drugs such as glucagon-like peptide-1 (GLP-1) receptor agonists (e.g., semaglutide, liraglutide, tirzepatide) reduced liver fat content by 5.2% and improved histological steatosis, hepatocellular ballooning, and lobular inflammation, with no significant effect on fibrosis (13). However, the underlying mechanisms are not fully understood.

Previous work in our laboratory identified *G protein coupled receptor* (GPCR) *kinase 5* (*Grk5*) as a candidate gene for adiposity in outbred rats, where *Grk5* expression in adipose tissue associates with increased adiposity (14). These results are supported in a global *Grk5* knockout (KO) mouse model where KO mice were protected against diet-induced adiposity/obesity along with decreased adipogenesis compared to wildtype (WT) littermate control mice (15). Interestingly, *GRK5* also associates with type 2 diabetes in humans (16). Our previous work and others have demonstrated that *Grk5* is highly expressed in stromal vascular cell fraction compared to mature adipocytes in visceral adipose tissues of mice and humans (17, 18, 19). Thus, we generated a *Grk5* knockout (KO) 3T3-L1 preadipocyte cell line, where KO cells displayed impaired adipocyte differentiation compared to WT cells, further demonstrating *Grk5* as an important gene for adiposity (17). GRK5 is a GPCR kinase that mediates downstream signaling through phosphorylation of activated receptors and recruitment of β-arrestin to facilitate receptor desensitization, internalization, and degradation (20). We found that deletion of GRK5 not only regulates GPCRs but also modulates Insulin-like growth factor 1 receptor (IGFR), a receptor tyrosine kinase, and its downstream Extracellular signal-regulated kinase 1/2 (ERK) signaling, resulting in impaired adipocyte differentiation (17). Since genetic modification is not a viable therapeutic option, it has sparked an increased interest in investigating potential GRK5 inhibitors as therapeutic candidates for chronic conditions (21, 22).

We have identified a small molecule GRK5 inhibitor, GRK5-IN-2, which preferentially inhibits GRK5 compared to other GRKs (17). *In vitro* treatment of WT 3T3-L1 preadipocytes with GRK5-IN-2 impaired both adipogenesis and lipogenesis compared to vehicle treated cells (17), suggesting a potential role for GRK5-IN-2 in modulating adipocyte biology and lipid metabolism *in vivo*. Thus, in this study, we aimed to determine whether GRK5 inhibition via GRK5-IN-2 *in vivo* could promote adipose tissue remodeling and improve metabolic health in diet-induced obesity.

## METHODS

### Animals

Mice were housed in standard cages under a 12-h light cycle and 12-h dark cycle (dark from 6:00 PM to 6:00 AM) at standard ambient temperature and humidity conditions and were provided with ad libitum water and a standard chow diet (Purina-LabDiet, Prolab RMH 3000). All experiments were performed using a protocol approved by the Institutional Animal Care and Use Committee at Wake Forest University School of Medicine in facilities approved by the American Association for Accreditation of Laboratory Animal Care.

Six weeks old male C57Bl/6J mice (Jackson Lab, Bar Harbor, ME, USA, Strain #000664) were used to assess the impact of GRK5-IN-2 (MedChem Express, Junction, NJ, USA, Cat#HY-136561) administration on diet-induced obesity and metabolic health. A treatment and follow-up study were conducted, each consisting of 10 mice per group and study designs can be found in **Fig. 1A** and **Fig. 4A**. Briefly, mice were fed a high fat diet (Research Diets Inc, New Brunswick, NJ, USA, D12451, 45% from fat) for 8 weeks to induce obesity.

**Figure 1.**
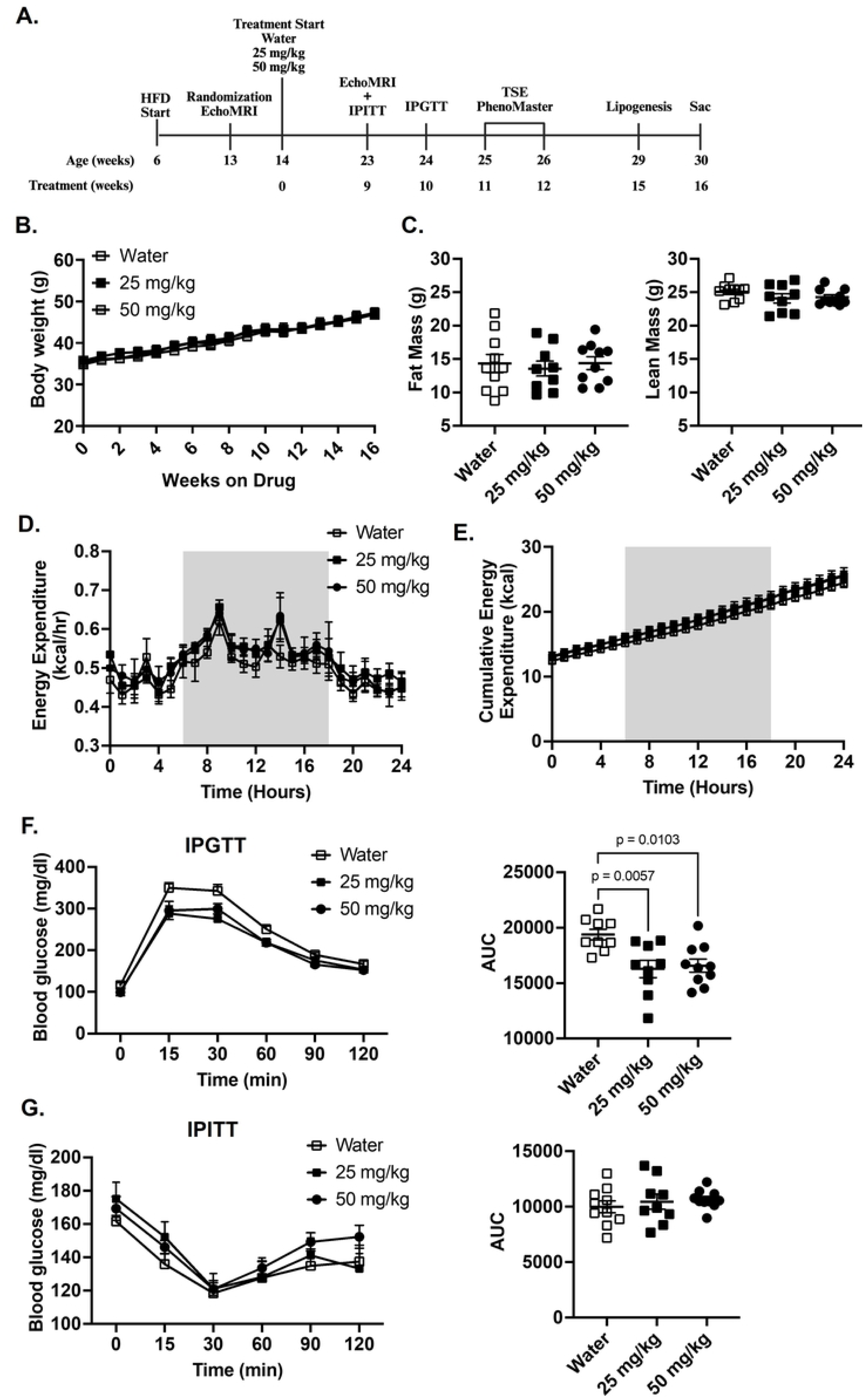
Effects of GRK5-IN-2 on adiposity and metabolic health in diet-induced obese mice. **(A)** Study design showing weeks of age and treatment. Six-week-old male C57Bl/6J mice were fed a high fat diet (HFD; 45% fat, D12451, Research Diets Inc) for 8 weeks to induce obesity prior to starting water control (open square), 25 mg/kg (closed square) or 50 mg/kg (closed circle) GRK5-IN-2 treatment. After 9 weeks of treatment, metabolic phenotyping started including EchoMRI, intraperitoneal insulin and glucose tolerance tests (IPITT and IPGTT), TSE PhenoMaster chambers, and a lipogenesis functional study. Mice were euthanized after 16 weeks of treatment (Sac). **(B)** Body weight was measured weekly (n=10/group). **(C)** Body composition was measured after 9 weeks of treatment (n=10/group). **(D-E)** After 11-12 weeks of treatment, mice (n=4/group) were used for indirect calorimetry (TSE PhenoMaster System). Results analyzed using the CalR Web Application (Version 1.3). Gray shading indicates a dark cycle. **(F-G)** Mice (n=10/group) were fasted for 16 and 4 hours in preparation for an IPGTT and IPITT, respectively. Blood glucose was measured at 0, 15, 30, 60, 90, and 120-minutes post injection. Area under the curve (AUC) was calculated to assess glucose and insulin sensitivity. All results are mean ± SEM. Statistical significance was assessed using one-way or two-way ANOVA.

Before treatment, body weight and fat mass were measured using EchoMRI™ (EchoMRI LLC, Houston, TX, USA) to randomize divided into groups: Control (water), 25 mg/kg, or 50 mg/kg dose of GRK5-IN-2 administered via oral gavage 5 days a week. Treatment lasted 16 weeks for the main study and 13 weeks for the follow-up study.

For both the treatment and follow-up study, body composition was measured using EchoMRI analysis after 9 weeks of drug treatment. Non-fasted mice were weighed and scanned using EchoMRI for 2 minutes in duplication.

After 9 and 10 weeks of treatment, mice went through intraperitoneal insulin and glucose tolerance tests (23), respectively. Briefly, mice were fasted for 4 hours or 16 hours prior to receiving an insulin (1 U/kg body weight, Eli Lilly, Indianapolis, IN, USA) or glucose (1 g/kg, Sigma-Aldrich, Milwaukee, WI, USA) injection, respectively. Blood glucose was measured (Contour Next EZ) from a tail snip at 0, 15, 30, 60, 90, and 120 minutes after injection. Insulin sensitivity and glucose tolerance was assessed by calculating the area under the curve (AUC).

After 11-12 weeks of treatment, mice spent 5 days single-housed in the TSE PhenoMaster environmental chambers (TSE Systems, Chesterfield, MO) to measure energy expenditure using indirect calorimetry where oxygen consumption, carbon dioxide production, respiratory exchange ratio (RER), food intake, energy expenditure, and locomotor activity were measured. RER near 0.7 indicates that fat is predominantly being used for energy, while RER near 1.0 indicates carbohydrates are predominant.

In the initial study, after 15 weeks of drug administration, a sub-set of treatment study mice (n=3/treatment group) were used for an *in vivo* lipogenesis analysis. Detailed procedures are shown below.

All other mice (n=7/treatment group) were euthanized, and serum was collected via cardiac puncture. Liver, gonadal visceral white adipose tissue (gWAT), inguinal subcutaneous (SubQ) WAT, retroperitoneal visceral white fat (RetroFat), brown adipose tissue (BAT), heart, kidney, and muscle were collected and flash frozen in liquid nitrogen. All samples were stored at -80°C until analysis.

### In Vivo Lipogenesis Analysis

Experimental non-fasted mice used for metabolic phenotyping (n=3/treatment) were used to assess *de novo* lipogenesis and fatty acid uptake and esterification using radiolabeled isotopes (23). Briefly, control and drug treated mice were injected with 5 μCi per mouse of [1,2-^14^C]-acetic acid (PerkinElmer, Waltham, MA, USA) or 50 μCi per mouse of [9,10-^3^H(N)]-oleic acid (PerkinElmer, Waltham, MA, USA) plus 0.04 mM oleic acid (Sigma-Aldrich) via retro-orbital injection. Tissues (Liver, gWAT, inguinal SubQ, BAT and muscle) were collected 3 hours after injection and snap frozen in liquid nitrogen until use. Lipids were extracted from tissues using hexanes:isopropanol (3:2, vol:vol) overnight. Lipid classes from standards and all tissues collected were separated by thin layer chromatography using Silica Gel plates and a solvent system containing hexane:diethyl ether:acetic acid (80:20:2, vol:vol:vol). Lipids were visualized by exposure to iodine vapor, and bands corresponding to triglycerides, free cholesterol, cholesteryl ester, and phospholipids were scraped and counted using a scintillation counter. Data was normalized to tissue weight.

### Adipose Tissue Histology

gWAT histology samples, from the follow-up study only, were fixed in 10% formalin for 1 week prior to being stored in 70% ethanol. Samples (n=10/treatment group) were sliced, mounted and stained with hematoxylin and eosin by the Comparative Pathology Core at Wake Forest University School of Medicine. Slides were imaged using the PANNORAMIC™ digital slide scanner (3DHISTECH, Budapest, Hungary), and adipocyte size was measured using Visiopharm (Hoersholm, Denmark). At least 5,000 adipocytes were measured per slide, and adipocyte size was averaged based on the individual animal. Adipocytes sized between 20 to 20,000 μm^2^ were included for analysis.

### Serum Analysis

Animal blood was collected via cardiac puncture, and serum was separated with centrifugation in 1800 xg for 18 minutes at 4°C and store at -80°C until analysis. Free fatty acid (FFA) concentration was measured in initial study mice (n=5-7/treatment) using a colorimetric assay (ZenBio, Durham, NC, USA, Cat#SFA-1). Triglycerides and total cholesterol levels (n=10/treatment) were measured in the follow-up study using the L-Type Triglyceride kit (FUJIFILM, Lexington, MA, USA) and the Cholesterol Reagent (Pointe Scientific, Inc., Canton, MI, USA). Alanine transaminase (ALT) and aspartate transaminase (AST) serum levels (n=10/treatment) were measured using the SimpleStep ELISA® kits (Abcam, Cambridge, UK, Cat#: ab282882 and ab263882, respectively) following manufacturer’s instructions.

### Tissue Triglyceride Analysis

Frozen liver and adipose tissues were weighed, and total lipids were extracted using hexane: isopropanol (3:2, vol:vol) incubated for 48 hours at room temperature. Lipid extracts were dried down at 65°C. Once dry, 1% Triton-X in chloroform was added to each sample and dried down again at 65°C. Samples were resuspended in ddH_2_O to yield the aqueous lipid extract for each sample. Triglyceride levels were measured using the L-Type Triglyceride kit (FUJUFILM) and normalized to the tissue wet weight.

### RNA Extraction and Real-time PCR

Total RNA was harvested from tissues using Qiazol Lysis Reagent and isolated by following the protocol described in the RNeasy Lipid Tissue Mini Kit (QIAGEN, Aarhus, Denmark). The concentration and quality of RNA were determined using a Nanodrop (ThermoFisher Scientific, Waltham, MA, USA) and standardized to 1 μg of RNA for cDNA synthesis. The cDNA was prepared with the High-Capacity cDNA Reverse Transcription Kit (ThermoFisher Scientific) and stored at -20°C until used for Real-Time PCR. Real-Time PCR was performed in duplicate on Applied Biosystems QuantStudio™ 3 Real-Time PCR System using Taqman® Fast Advanced Master Mix and TaqMan® gene expression assays (ThermoFisher Scientific) including *Grk5* (Mm00517039_m1), *Peroxisome proliferator-activated receptor gamma* (*Pparγ*) (Mm0040940_m1), *Adiponectin* (Mm04933656_m1), *CCAAT/enhancer-binding protein alpha* (*Cebpα*) (Mm00514283_s1), *Fatty acid binding protein 4* (*Fabp4*) (Mm00445878_m1), *Fatty acid synthase* (*Fasn*) (Mm00662319_m1), *Chemokine ligand 1* (*Cxcl1*) (Mm04207460_m1), *C-X-C motif chemokine ligand 2* (*Cxcl2*) (Mm00436450_m1), *Peroxisome proliferator-activated receptor alpha* (*Pparα)* (Mm00440939_m1), *Sterol regulatory element-binding transcription factor* 1 (*Srebf-1*) (Mm00550338_m1), *Peroxisome proliferator-activated receptor gamma coactivator-1-alpha* (*Pgc-1α*) (Mm01208835_m1), *Interleukin* 6 (*Il6*) (Mm00446190_m1), *Collagen, type VI, alpha 3* (*Col6a3*) (Mm00711678_m1), *Acetyl-CoA carboxylase 2 (Acc2*) (Mm01204671_m1), and *Mitochondrial transcription factor A (Tfam)* (Mm00447485_m1). Gene expression was normalized to the *18s* rRNA (REF4354655) endogenous control and analyzed using the 2ddct method with 95% confidence.

### Protein Extraction and Western blot

Total cellular protein from liver was harvested in Pierce™ IP lysis buffer (ThermoFisher Scientific, Cat#87787) supplemented with cOmplete™ EDTA-free protease (Sigma-Aldrich, Cat#11873580001) and PhosSTOP™ Phosphatase (Sigma-Aldrich, Cat#4906845001) inhibitor tablets and frozen at -20°C until used. Protein samples were normalized to 1 mg of protein, prepared in non-reducing laemmli buffer and DTT, and heated at 95°C for 10 minutes. Protein was loaded and separated on a 4-20% polyacrylamide gel (Bio-Rad Laboratories, Hercules, CA, USA, Cat#5671094) and transferred to a 0.2 μm nitrocellulose membrane (Bio-Rad, Cat#1620112). Membranes were blocked in 5% non-fat milk in 1X Tris-buffered saline plus 0.1% Tween (TBST, Bio-Rad) for 2 hours at room temperature. Primary antibodies were diluted in TBST with 1% non-fat dry milk and incubated overnight at 4°C with gentle rocking. Primary antibodies were diluted as follows: α-tubulin (Cell Signaling Technology, #2144) at 1:1000, Cytochrome c oxidase subunit 4 (COXIV) (Cell Signaling Technology, #11967) at 1:1000, and Acyl-CoA synthetase long-chain family member 1 (ACSL1) (Cell Signaling Technology, #9189) at 1:1000. Following overnight incubation, membranes were washed 3 times in TBST for 5 minutes with agitation and incubated with secondary antibody in 5% non-fat milk for 1 hour at room temperature (ThermoFisher Scientific mouse and rabbit secondaries, 1:5000) with gentle rocking. Membranes were washed 3 times in TBST for 5 minutes with agitation. SuperSignal™ West Pico PLUS Chemiluminescent Substrate (ThermoFisher Scientific, Cat#24580X4) was added to the membrane prior to imaging using the ChemiDoc Gel Imaging System (Bio-Rad). Protein expressions were quantified using Bio-Rad ImageLab software.

### Statistics

Data are presented as mean ± standard error of the mean (SEM) throughout the figures. In the initial treatment study, binary comparisons were performed using one-way ANOVA. Since the follow-up study focused on only the 25 mg/kg dose, binary comparisons were performed using a two-tailed Student’s unpaired *t* test. Body weight and systemic glucose and insulin tolerance tests were analyzed using a two-way ANOVA with repeated measures. Prism 10 software (GraphPad) was used to perform statistical analyses (Statistical significance *p* < 0.05). CalR, a web-based tool, was used to perform statistical ANCOVA analyses (Statistical significance *p* < 0.05) for indirect calorimetry measurements. All graphs were generated using Prism 10 software.

## RESULTS

### GRK5-IN-2 treatment did not affect whole-body or adipose tissue characteristics or function in diet-induced obese mice

Previous studies have reported that deletion of *Grk5* results in decreased adipocyte differentiation (17) and protection against diet-induced adiposity (15), supporting GRK5 signaling as a therapeutic target for obesity. Additionally, Amlexanox, a non-specific GRK inhibitor (24), has been shown to prevent and reverse diet-induced obesity in mice at oral doses ranging from 25-100 mg/kg (25). Based on these findings, we hypothesized that inhibition of GRK5 using GRK5-IN-2 would reduce adiposity and improve metabolic health in a diet-induced obese mouse model. To test this, we orally treated diet-induced obese mice with either water (control) or GRK5-IN-2 (25 mg/kg or 50 mg/kg) and performed a series of metabolic phenotyping experiments (**Fig. 1A**). We found no significant differences in body weight or composition between control and treatment groups (**Fig. 1B-C**). Similarly, there were no significant differences in energy expenditure (**Fig. 1D**), cumulative food intake (**Fig. 1E**), or other indirect calorimetry parameters (**Supplemental Fig. 1A**).

After 10 weeks of drug administration, mice treated with either 25 mg /kg and 50 mg/kg doses of GRK5-IN-2 showed significant improvement in glucose tolerance compared to water-treated controls (**Fig. 1F**, p=0.0057 and p=0.0103, respectively), with no differences between the two doses. No changes were observed with insulin sensitivity between treatment groups (**Fig. 1G**). At the end of the study, consistent with the body composition results, no significant differences were observed in the wet weights of gWAT, SubQ, RetroFat, or BAT between treatment groups (**Supplemental Fig. 1B**). Fasting serum was collected to assess lipid homeostasis parameters, but no differences in serum free fatty acids were detected with either dose compared to water-treated mice (**Supplemental Fig. 1C**).

To further assess the functional impact of GRK5-IN-2 on adipose lipid metabolism, we conducted a tracer-based study using [¹⁴C]-acetic acid and [³H]-oleic acid to evaluate de novo lipogenesis and fatty acid uptake and incorporation into triglycerides. We found that there were no statistically significant differences in triglyceride synthesis rates between treatment groups (**Fig. 2A**). No changes were observed in the synthesis of cholesteryl esters, phospholipids, or free cholesterol from [^14^C]-acetic acid (**Fig. 2B**). Similarly, GRK5 inhibition did not significantly affect [^3^H]-oleic acid uptake (**Fig. 2C**) or its incorporation into triglycerides, cholesteryl ester, or phospholipids (**Fig. 2D**). Together, these data indicate that GRK5-IN-2 does not substantially alter systemic and adipose tissue metabolism in diet-induced obese mice.

**Figure 2.**
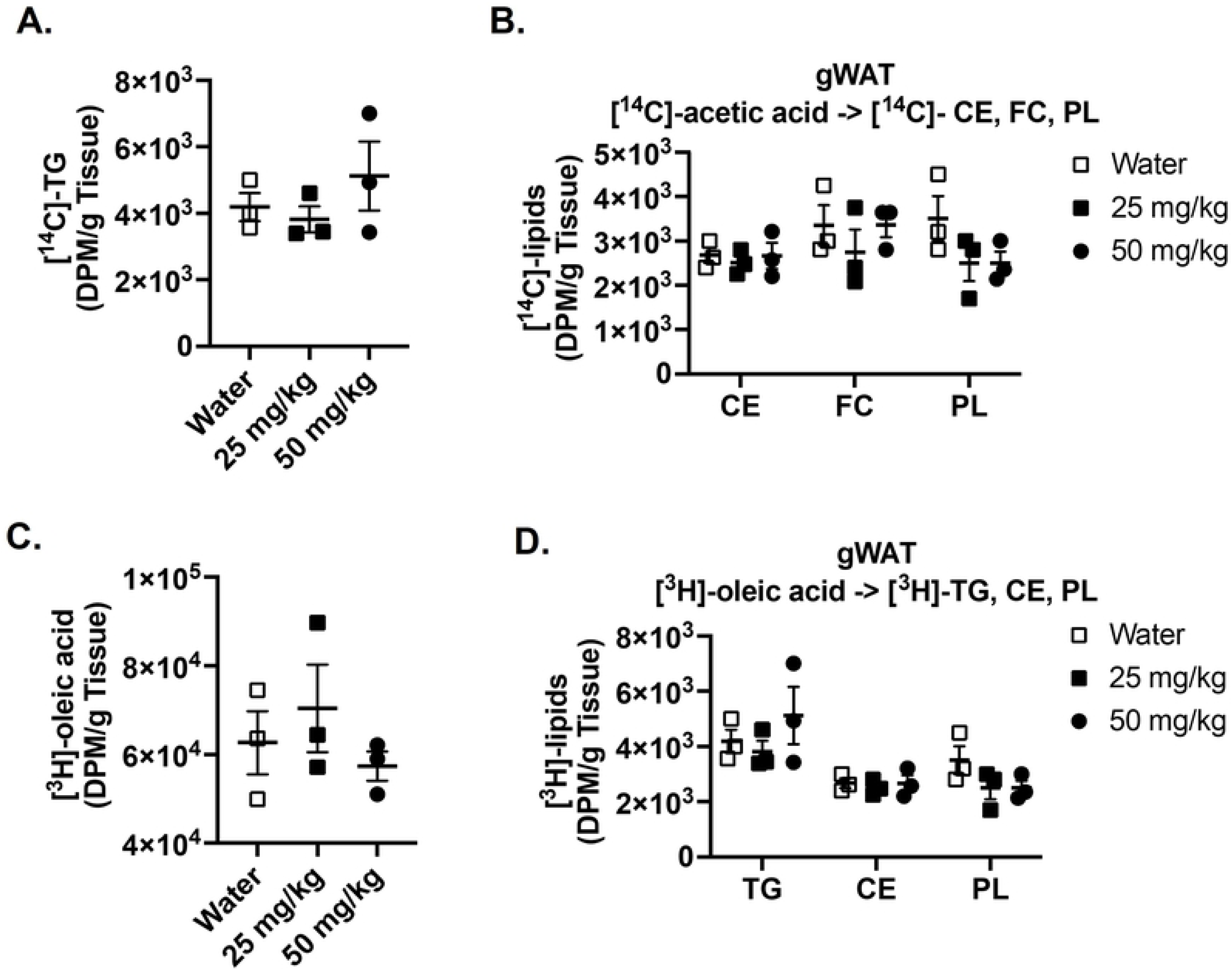
Effects of GRK5-IN-2 on visceral adipose tissue functionality in diet-induced obese mice. *De novo* and fatty acid uptake were measured using radiolabeled [^14^C]-acetate and [^3^H]-oleate. Control and GRK5-IN-2 treated mice (n=3/group) were injected with 5 μCi per mouse of [1,2-^14^C]-acetic acid (**A-B**) or 50 μCi per mouse of [9,10-^3^H(N)]-oleic acid (**C-D**). Tissues were collected for 3 hours post injection. Lipids were extracted and separated using thin layer chromatography. [^14^C]-triglycerides (TG), [^14^C]-cholesteryl esters (CE), [^14^C]-free cholesterols (FC), [^14^C]-phospholipids (PL), [^3^H]-oleic acids, [^3^H]-TG, [^3^H]-CE, and [^3^H]-PL were quantified by liquid scintillation counting. All results are mean ± SEM. Statistical significance was assessed using one-way ANOVA.

### GRK5-IN-2 treatment decreased diet-indued liver steatosis compared to control mice

Previous studies using whole-body *Grk5* knockout mice reported the development of severe hepatic steatosis compared to WT controls (26). In contrast, treatment with Amlexanox, a non-specific GRK inhibitor, was shown to ameliorate hepatic steatosis (25). This conflicting evidence prompted us to examine the effects of GRK5-IN-2, a preferred GRK5 inhibitor, on hepatic lipid metabolism. There were no significant differences in liver weight or liver-to-body weight ratio between control and treatment groups (**Fig 3A-B**). Interestingly, GRK5-IN-2 (both 25 and 50 mg/kg) significantly reduced total hepatic triglyceride content relative to water-treated control obese mice (**Fig 3C**, p<0.0001).

**Figure 3.**
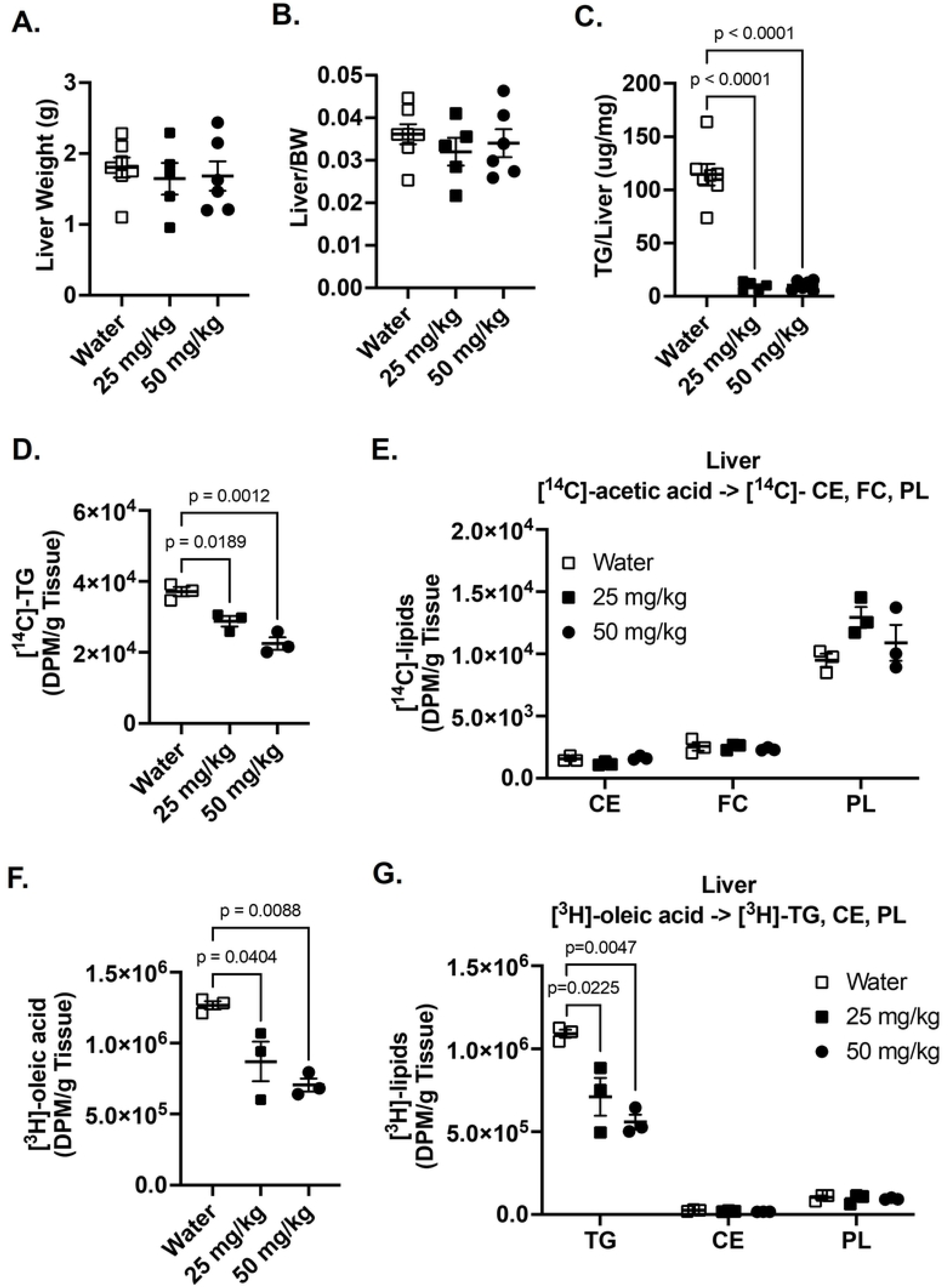
Effects of GRK5-IN-2 on liver lipid metabolism in diet-induced obese mice. **(A-B)** At study end point, mouse liver (n= 5-7/group) was collected, weighed, and normalized to total body weight. **(C)** Liver lipid content was extracted (n=15/group), and triglyceride content was measured using a colorimetric assay normalized to liver tissue weight. After 15 weeks of GRK5-IN-2 treatment, *de novo* lipogenesis and fatty acid uptake were measured using radiolabeled isotopes. Control and drug treated mice (n=3/group) were injected with 5 μCi per mouse of [1,2-^14^C]-acetic acid **(D-E)** and 50 μCi per mouse of [9,10-^3^H(N)]-oleic acid **(F-G)**. Tissues were collected for 3 hours post injection. Lipids were extracted and separated using thin layer chromatography. [^14^C]-triglycerides (TG), [^14^C]-cholesteryl esters (CE), [^14^C]-free cholesterols (FC), [^14^C]-phospholipids (PL), [^3^H]-oleic acids, [^3^H]-TG, [^3^H]-CE, and [^3^H]-PL were quantified by liquid scintillation counting. All results are mean ± SEM. Statistical significance was assessed using one-way ANOVA.

We further evaluated functional lipid synthesis in liver by performing radiolabeled tracer assays using [¹⁴C]-acetic acid and [³H]-oleic acid. Both GRK5-IN-2 doses significantly decreased *de novo* biosynthesis of triglycerides from [^14^C]-acetic acid relative to control (**Fig. 3D**, p=0.0189 and p=0.0012), without affecting the synthesis of other lipid species (**Fig. 3E**). GRK5-IN-2 treatment also significantly decreased [^3^H]-oleic acid uptake (**Fig. 3F**, p=0.0404 and p=0.0088) and reduced fatty acid incorporation into triglycerides (**Fig. 3G**, p=0.0225 and p=0.0047) but had no impact on the synthesis of other lipids **(Fig. 3G)**. These data suggest that GRK5-IN-2 treatment improves hepatic steatosis in diet-induced obese mice, potentially through suppression of fatty acid uptake and triglyceride biosynthesis.

### A follow-up study supports that GRK5-IN-2 treatment reduces hepatic steatosis with minimal effects on adiposity compared to water vehicle-treated mice in obesity

To further investigate downstream molecular changes of GRK5-IN-2 treatment, we conducted a follow-up study focusing on the 25 mg/kg dose (**Fig. 4A**), as no significant differences were observed between 25 mg/kg and 50 mg/kg doses in the initial study. In this follow-up study, consistent with our prior findings, no differences were observed in body weight (**Fig. 4B**), body fat/lean composition (**Fig. 4C**), adipocyte size or number in adipose tissue (**Fig. 4D**), or adipose triglyceride content (**Fig. 4E**) between treatments. GRK5-IN-2 treatment also did not significantly alter indirect calorimetry parameters (**Data not shown**), circulating lipid profiles (**Supplementary Fig. 2A**), or insulin tolerance test **(Supplementary Fig. 2B)**. However, in contrast to our initial findings, GRK5-IN-2 treatment did not improve glucose tolerance test relative to water-treated obese controls in this follow-up cohort **(Supplemental Fig. 2C**).

**Figure 4.**
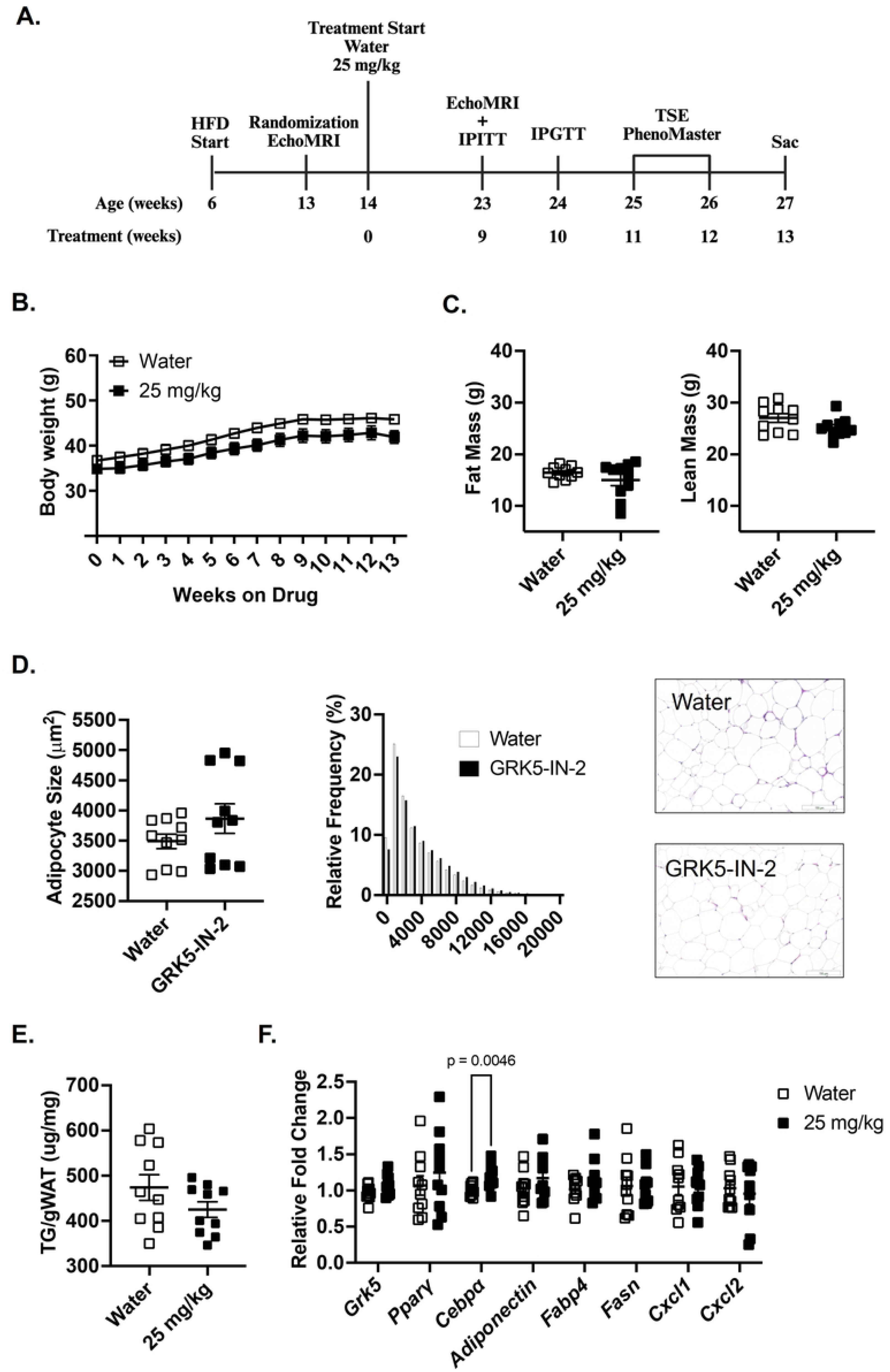
Effects of GRK5-IN-2 treatment on adipose and systemic metabolism in a follow-up study. **(A)** Study design illustrating timeline and treatments. Six-week-old male C57Bl/6J mice were fed a high-fat diet (HFD; 45% fat, D12451, Research Diets Inc.) for 8 weeks to induce obesity before initiating GRK5-IN-2 treatment. After 9 weeks of treatment, metabolic phenotyping was conducted, including EchoMRI, intraperitoneal insulin (IPITT) and glucose tolerance tests (IPGTT), TSE PhenoMaster metabolic chambers, and a lipogenesis functional assay using radiolabeled tracer. Mice were euthanized after 13 weeks of treatment (Sac). **(B)** Body weight measured weekly (n=10/group). **(C)** Body composition assessed via EchoMRI after 9 weeks of treatment (n=10/group). **(D)** Gonadal white adipose tissue (gWAT) was collected, fixed in 10% formalin, and stained with hematoxylin and eosin (H&E) for histological analysis (n=10/group). Representative H&E-stained gWAT images from water- and GRK5-IN-2-treated mice. **(E)** Triglyceride (TG) content in gWAT lipids extracted and quantified using a colorimetric assay (n=10/group). **(F)** RNA was extracted from gWAT (n=10/group), reverse-transcribed to cDNA, and analyzed via real-time PCR to quantify adipogenic and inflammatory gene expression normalized to 18S rRNA (endogenous control). Results are expressed as fold change relative to the water-treated group. All data are presented as mean ± SEM. Statistical significance was assessed using two-tailed Student’s unpaired *t* test.

To investigate if there are molecular changes in adipose tissue, we quantified the expression of genes involved in adipogenesis (Pparγ, Cebpα), lipogenesis (Fasn), adipocyte markers (Adiponectin, Fabp4)(27). We also examined gene expressions of Grk5 and inflammatory markers (Cxcl1, Cxcl2)(28) in gWAT from diet-induced obese mice treated with water (vehicle control) or GRK5-IN-2. Relative to controls, no differences were observed in Grk5, Pparγ, Fasn, Adiponectin, Fabp4, Cxcl1, or Cxcl2 expression, except for a significant change in Cebpα expression (**Fig. 4G**, p=0.0046). Together, these data support our previous findings that GRK5-IN-2 treatment has minimal effects on adipose tissue in diet-induced obese mice.

To assess the effects of GRK5-IN-2 on liver biology in the follow-up cohort of diet-induced obese mice, we measured hepatic triglyceride content and serum markers of liver function. Consistent with initial findings, GRK5-IN-2 treatment significantly reduced total hepatic triglyceride content which corresponded to a reduction in percent liver triglyceride levels compared to water-treated controls (**Fig. 5A**, p=0.0197). No liver toxicity was observed, as indicated by no differences in the serum ALT/AST ratio between treatment groups (**Fig. 5B**).

**Figure 5.**
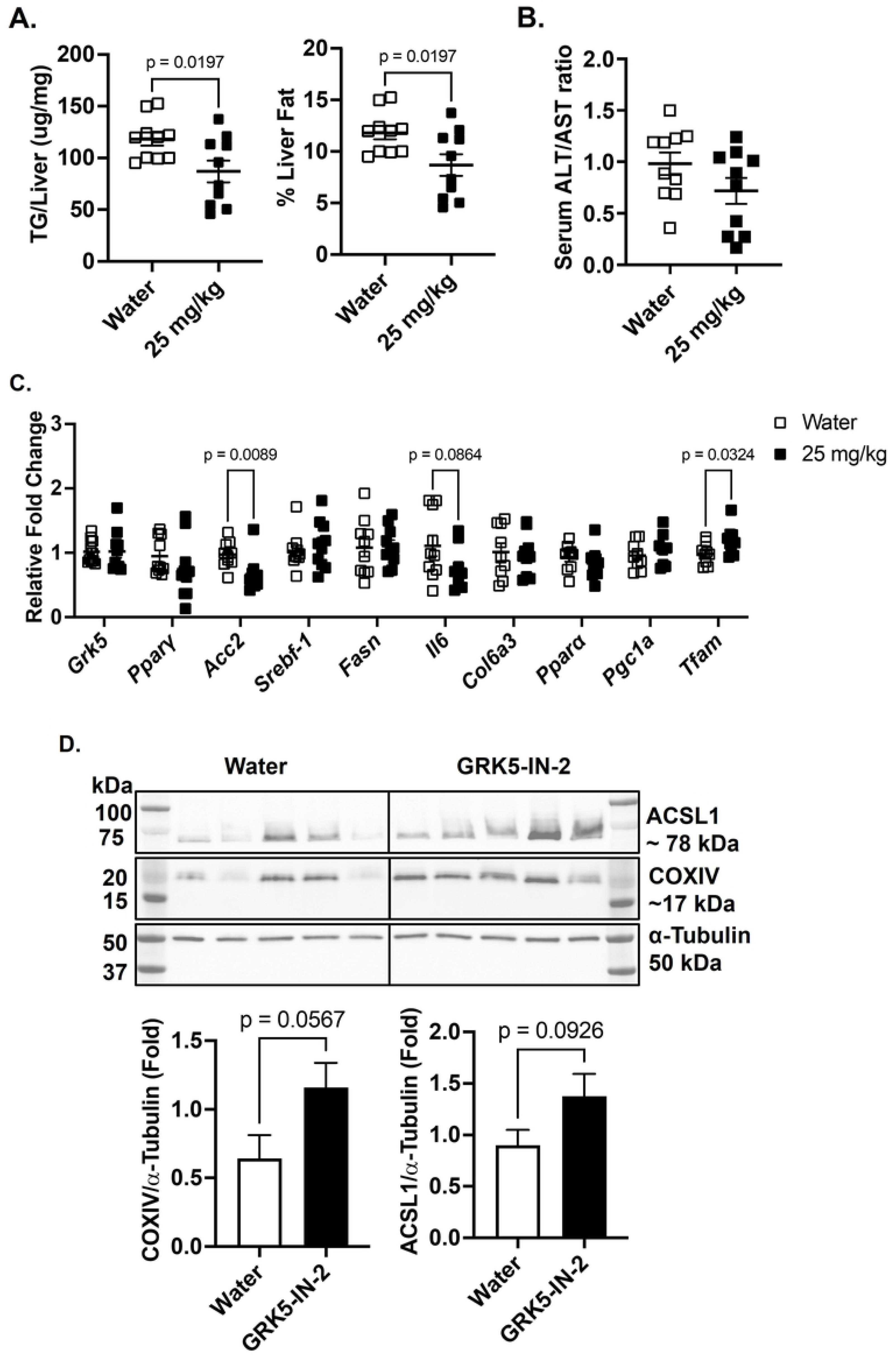
Effects of GRK5-IN-2 on liver triglyceride content, function, and molecular profiles in a follow-up study. **(A)** Hepatic lipids were extracted from liver tissue (n=10/group), and total triglyceride (TG) content and percent liver TG levels were quantified using a colorimetric assay. **(B)** Serum alanine aminotransferase (ALT) and aspartate aminotransferase (AST) levels were measured using a colorimetric assay to calculate the ALT/AST ratio (n=10/group). **(C)** RNA was extracted from liver tissue (n=10/group), reverse-transcribed to cDNA, and analyzed via real-time PCR to quantify expression of genes involved in lipogenesis, inflammation, fibrosis, fatty acid oxidation, and mitochondrial biogenesis, normalized to 18S rRNA (endogenous control). Results are expressed as fold change relative to the water-treated group. **(D)** Liver protein (n=10/group) was extracted and subject to Western blot using anti-ACSL1, anti-COXIV, and anti-α-Tubulin antibodies. All data are presented as mean ± SEM. Statistical significance was assessed using two-tailed Student’s unpaired *t* test.

To further investigate the molecular effects of GRK5-IN-2 on liver biology in diet-induced obese mice, we analyzed gene expression in liver tissue, focusing on lipogenesis (*Pparg*, *Srebf1*, *Acc2*, *Fasn*), inflammation (*Il6*), and fibrosis (*Col6a3*). GRK5-IN-2 treatment significantly suppressed *Acc2* expression (*p*=0.0089) compared to water-treated controls, but did not alter *Grk5*, *Pparg*, *Srebf1*, or *Fasn* expression (**Fig. 5C**). A trend toward reduced *Il6* expression (*p*=0.0864) was observed, with no difference in *Col6a3* expression (**Fig. 5C**).

We also examined genes involved in fatty acid oxidation (*Ppara*, *Ppargc1a*) and mitochondrial biogenesis (*Tfam*). While *Ppara* and *Ppargc1a* expression remained unchanged, GRK5-IN-2 treatment significantly increased *Tfam* expression (**Fig. 5C**, *p*=0.0324). Additionally, Western blot analysis revealed trends toward increased protein expression of COXIV (*p*=0.0567), a subunit of mitochondrial complex IV, and ACSL1 (*p*=0.0926), an enzyme converting long-chain fatty acids to fatty acyl-CoA (**Fig. 5D**). These increases in COXIV and ACSL1 suggest enhanced mitochondrial oxidative capacity, potentially promoting fatty acid oxidation (29, 30). Collectively, these findings indicate that GRK5-IN-2 may mitigate liver steatosis by favoring fatty acid utilization over triglyceride synthesis.

## DISCUSSION

In the present study, we examined the therapeutic effects of a GRK inhibitor, GRK5-IN-2, on adiposity, hepatic steatosis, and systemic metabolism in diet-induced obese mice. Although the GRK5 inhibition does not affect adiposity or changes in energy expenditure in the initial (**Fig. 1**, **Fig. 2, and Supplemental Fig. 1**) and follow-up studies (**Fig. 4 and Supplemental Fig. 2)**, it consistently reduced hepatic triglyceride content in both studies (**Fig. 3 and Fig. 5**). Additionally, in the follow-up study, GRK5-IN-2 treatment significantly suppressed lipogenic gene expression, particularly Acc2, and enhanced mitochondrial biogenesis gene expression, *Tfam*, in the liver (**Fig. 5**). GRK5-IN-2 treatment also showed trends toward increased protein expression of fatty acid oxidation markers COXIV and ACSL1 in the liver (**Fig. 5**). These findings suggest that GRK5-IN-2 treatment may alleviate hepatic steatosis by modulating both liver lipid anabolism and catabolism pathways.

In the initial cohort, GRK5-IN-2 at 25 mg/kg and 50 mg/kg improved glucose tolerance without altering adiposity compared to water-treated controls (**Fig. 1**). As no dose-dependent differences were observed in metabolic parameters, the follow-up study used the 25 mg/kg dose. However, the glucose tolerance improvement was not replicated (**Fig. 4**), suggesting that the metabolic benefit may not be robust. This variability is consistent with human trials of Amlexanox, where glucose control improved only in patients with elevated baseline inflammation (e.g., high C-reactive protein), highlighting cohort-specific responses(31). In mice, discrepancies in glucose tolerance can arise from cohort-specific factors, including differences in fat and lean mass, which significantly influence insulin sensitivity and glucose uptake (32, 33). Lastly, variations in dosing protocols or environmental conditions (e.g., housing, handling stress) may also contribute to the inconsistent IPGTT results (34).

Our findings also differ from the whole body *Grk5* KO mouse model, which protected against diet-induced obesity (15). A key difference lies in tissue specificity and developmental timing. GRK5 is more highly expressed in preadipocytes compared to mature adipocytes (17), and the *Grk5* KO model disrupts early adipogenesis during development. In adult mice, preadipocytes constitute approximately 5-20% of adipose tissue cells (35), limiting the potential impact of GRK5-IN-2 on adipogenesis. Our results also differ from mice treated with Amlexanox, which reduced fat mass and improved metabolic health (25). Amlexanox, a non-specific GRK inhibitor (24), also inhibits TBK1 and IKKε, reducing adipose inflammation. Since chronic inflammation contributes to adipose tissue dysfunction and obesity, the metabolic improvements observed with Amlexanox may result from its anti-inflammatory effects (36, 37). Thus, the lack of effect of GRK5-IN-2 on adipose inflammatory genes distinguishes it from Amlexanox, which exerts anti-inflammatory effects and attenuates diet-induced obesity.

Although we did not see differences in adipose tissue biology, we found that diet-induced obese mice treated with GRK5-IN-2 had decreased diet-induced liver triglyceride accumulation compared to controls in both studies (**Fig. 3** and **Fig. 5**). Hepatic steatosis is a defining characteristic of MASLD (10), which is strongly associated with diet-induced obesity. Our findings are similar to previous work showing that Amlexanox, a non-specific GRK inhibitor, protected mice against diet-induced obesity and liver steatosis (24, 25). However, these results contrast with the whole body *Grk5* KO mouse model, which found increased hepatic steatosis relative to controls (26). This discrepancy may be due to differences in how each approach affects hepatic insulin sensitivity. The whole body *Grk5* KO mice exhibit impaired insulin signaling, which contributes to severe hepatic steatosis (26). In contrast, Amlexanox is not only a non-specific GRK inhibitor but also a dual inhibitor of TBK1/IKKε, which reduces inflammation and inflammation-related insulin resistance, resulting in improved liver steatosis (24, 25).

GRK5-IN-2 reduces liver triglyceride content in diet-induced obese mice by decreasing lipogenesis (**Fig. 3**) and increasing mitochondrial fatty acid oxidation (**Fig. 5**), though its precise mechanisms remain unclear. Despite preferentially inhibiting GRK5, GRK5-IN-2 also binds GRK2 (17), which is highly expressed in hepatocytes (38), the primary cells driving liver lipid metabolism (39), unlike the low-expressed GRK5 (40, 41). GRK2 is known to translocate to mitochondria in stress models, such as heart failure, where its reduction enhances ATP production and fatty acid utilization (42). Thus, GRK5-IN-2 may inhibit GRK2 translocation to liver mitochondria, promoting lipid catabolism and mitochondrial function. We hypothesize that GRK5-IN-2 likely reduces hepatic lipid accumulation by suppressing triglyceride synthesis and enhancing fatty acid oxidation, potentially via GRK2 inhibition, however, further studies are needed to confirm these mechanisms.

In summary, GRK5-IN-2, a small molecule GRK5 inhibitor, did not mitigate diet-induced obesity or improve adipose tissue health but significantly reduced hepatic lipid accumulation. Our findings suggest that GRK5-IN-2 attenuates hepatic steatosis by suppressing triglyceride synthesis and promoting mitochondrial fatty acid oxidation, potentially through inhibition of GRK2 alongside GRK5. These findings underscore the therapeutic potential of targeting GRK signaling to manage diet-induced hepatic steatosis.

## Acknowledgments

The authors wish to acknowledge services provided by the Comparative Pathology Laboratory Core (Department of Pathology/Section on Comparative Medicine) and the Lipid, Lipoprotein and Atherosclerosis Analysis Laboratory (Department of Internal Medicine/Section on Molecular Medicine) of the Wake Forest School of Medicine. The authors also wish to acknowledge the support of the Atrium Health Wake Forest Baptist Comprehensive Cancer Center Proteomics and Metabolomics Shared Resource, supported by the National Cancer Institute’s Cancer Center Support Grant award number P30CA012197. The content is solely the responsibility of the authors and does not necessarily represent the official views of the National Institutes of Health.

## Author contributions

MES: Designed the study, acquired data, analyzed and interpreted data, wrote the original draft. SS: Acquired data. TER: Interpreted data. CCK and LSW: Designed the study, interpreted data, revised the manuscript, acquired funding, and approved the final version for publication.

